# Microsurgical Isolation and Molecular Characterization of the Mouse Left Internal Mammary Artery: Insights into Natural Resistance to Atherosclerosis

**DOI:** 10.64898/2026.01.08.698440

**Authors:** Hebaallaha Hussein, Peter Saad, Kenneth W. Jenkins, Kristoforos Proko, Louis Zheng, Sowmitri Karthikeya Mantrala, Bijinu Balakrishnan, Manoj Bhasin, Vishwajeet Puri

## Abstract

Atherosclerosis develops unevenly across the vascular tree, yet the molecular basis for this regional susceptibility remains poorly defined. The left internal mammary artery (LIMA), the most durable conduit for coronary artery bypass (CABG) surgery, is uniquely resistant to atherosclerosis in humans; however, it has never been isolated or studied in mouse models, limiting mechanistic insight into atheroprotective pathways. Here, we establish the first method to identify and isolate the murine LIMA and generate the first transcriptomic atlas of this artery. Cross-species analyses of human and mouse LIMA reveal a conserved protective signature that distinguishes the LIMA from atheroprone vessels and uncover fundamental molecular differences that govern vascular resilience. This study presents a transformative experimental platform for dissecting the determinants of atheroprotection and identifying molecular targets to improve CABG graft performance and long-term cardiovascular outcomes.

## INTRODUCTION

Ischemic heart disease is the leading cause of death in the United States [1–3]. Various treatments have been developed to combat atherosclerotic disease formation; however, Coronary Artery Bypass Grafting (CABG) has been shown to provide the most significant survival advantage in patients with multivessel coronary artery disease, especially those with three-vessel involvement or left main coronary artery disease [1–6]. CABG significantly limits the new infarcts and reduces the procedural mortality and morbidity by providing flow distal to vessel occlusions. CABG surgery uses a conduit such as the saphenous vein, radial artery, or internal mammary artery (IMA) to bypass the occluded vessels and restore blood flow to the infarcted myocardium [2, 7–9].

The IMA, Internal Mammary Artery, commonly called the internal thoracic artery, is resistant to atherogenic effects of systemic cardiovascular risk factors, such as high LDL-C, diabetes, and hypertension [10]. Irrespective of their systemic nature, the IMA rarely develops atherosclerotic plaques with prevalence rates of histologically proven lesions ranging between 3.1% and 4.2% in unselected individuals [11, 12] and 0.7% and 7% in patients with multivessel coronary artery disease [13, 14]. Because of these reasons, Left IMA (LIMA) is the most common conduit used in CABG and has been considered the “gold standard” conduit for coronary artery bypass grafting despite evidence that the right mammary artery is identical [15]. Although the saphenous vein graft is also used, its patency in a 10-year angiography follow-up is only 61%, whereas for LIMA it is 85% [16, 17]. LIMA also matches in size with the coronary artery, the most important coronary artery in the human circulation, the LAD, which, cumulatively, also makes it an adequate conduit over the saphenous vein graft [18, 19]. In addition to revascularizing coronary lesions, LIMA has more advantages. For example, each bypass protects the native coronary artery from existing and subsequent coronary disease. Postmortem microscopic examinations demonstrated that LIMA was dramatically protected against atherosclerosis [11, 20].

Interestingly, LIMA’s remarkable resistance to atherosclerosis in humans is proposed to be attributed in part to its unique endothelial features. Its intimal layer is composed of distinct, regularly organized endothelial cells with low intercellular permeability, reducing endothelial injury and limiting immune-cell trafficking. Also, LIMA’s from CABG patients show minimal endothelial denudation, and their vasodilatory function remains well preserved compared to athero-prone arteries. Also, LIMA is less affected by aging-associated endothelial dysfunction, owing to higher activity of antioxidant defense enzymes such as SOD relative to carotid arteries [10, 21]. Furthermore, LIMA has fewer endothelial fenestrations, decreased intracellular junction permeability, and production of key mediators such as nitric oxide and prostacyclins, which reduce inflammation and plaque development [2]. Proteomic studies revealed that the intimal layer of human LIMA significantly expresses fewer proteoglycans, which are the main components, along with VSMCs, of advanced atheroma [10].

The unique long-term patency and atheroresistant properties of the LIMA make it an important model for uncovering the molecular mechanisms underlying vascular protection and for identifying novel therapeutic strategies against atherosclerosis. Progress in this area, however, has been hindered by the limited availability of human LIMA tissue, restricting mechanistic investigations. Because the vascular branching pattern of the aortic arch is highly conserved between mice and humans [22], murine systems offer a viable and powerful platform for studying human vascular disease. Yet, despite the widespread use of mouse models to investigate atherogenesis, the murine LIMA has never been isolated or characterized, representing a major gap in the field. In the present study, we address this gap by establishing a novel microsurgical protocol for isolating the mouse LIMA, thereby enabling detailed mechanistic studies of its intrinsic anti-atherosclerotic properties in a genetically tractable model system. We further perform comprehensive molecular characterization of the LIMA and aorta in atherosclerosis-prone mouse models and compare these signatures to human LIMA and athero-susceptible arteries, strengthening the translational relevance and impact of our approach.

## MATERIALS & METHODS

### MATERIALS

Mice and CO_2_ supply machine (CO_2_ induction system); 25 G needles (or pins or adhesive tape), 10 ml syringes and 1.5 ml Eppendorf tubes; Blood collection tubes (BD® 365967 Microtainer®); 70 % EtOH and Phosphate-buffered saline (PBS); TRIzol (Thermo Fisher Scientific, catalog number: 15596026; Direct-zol^TM^ RNA MicroPrep W/Tri Reagent (ZYMO RESEARCH, catalog number: R2061-A; Dissection scissors (Fine Science Tools, catalog number: 91460-11); Fine iris scissors (Fine Science Tools, catalog number: 14094-11); Spring scissors (Fine Science Tools, catalog number: 15009-08); Curved forceps (Fine Science Tools, catalog number: 11073-10); Dissecting Microscope with camera (e.g., Motic SMZ-171 BP-stereo microscope).

### METHODS

#### Mice

Homozygous Apoe-knockout (Apoe-/-) mice were bought from Jackson lab. The experimental protocols were conducted in accordance with the Institutional Animal Care and Use Committee (IACUC) approval (IACUC Protocol # 00000030) guidelines. Ohio University holds an Animal Welfare Assurance (A3610-01) from the NIH-Office of Laboratory Animal Welfare and complies with the USDA Animal Welfare Act Regulations under license 31-R-0082. Care and use of laboratory animals adhered to the *Guide for the Care and Use of Laboratory Animals.* The mouse colonies were maintained in racked individual ventilation cages according to current national legislation. The mice had dust and pathogen-free bedding and sufficient nesting and environmental enrichment material for the development of species-specific behavior. All mice had ad libitum access to food and water in environmental conditions of 45–65% relative humidity, temperatures of 21–24 °C and a 12–12 h light–dark cycle.

#### Aorta isolation

For aorta isolation, mice were maintained in individually ventilated cages under national regulatory standards, with dust- and pathogen-free bedding, ample nesting and enrichment materials, ad libitum access to food and water, 45–65% humidity, temperatures of 21–24 °C, and a 12:12-h light–dark cycle. Clean dissection instruments were prepared prior to the procedure. Mice were euthanized by CO inhalation in an IACUC-approved chamber, positioned supine on a Styrofoam surface, and secured with surgical tape. The ventral abdominal fur was saturated with 70% ethanol to prevent contamination, and a small incision was made in the lower abdomen to initiate dissection. The skin was lifted with forceps and separated from the abdominal wall, after which the abdominal wall was opened along the Linea alba to the level of the ribcage, exposing the xiphoid process. The xiphoid was grasped and lateral incisions were made beneath the ribcage to expose the diaphragm, which was then incised along its length to open the pleural cavity. The ribs lateral to the mediastinum were carefully cut with curved blunt scissors to expose the heart while preserving the internal mammary arteries. Residual blood was removed with sterile gauze, and the lungs were excised to fully expose the heart and aorta. For perfusion, 20 mL of sterile ice-cold PBS was gently infused into the left ventricle using a 10-cc syringe fitted with a 25-gauge needle; the right atrium was incised to relieve pressure, and perfusate was absorbed with sterile gauze. Following perfusion, excess fluid was removed, and abdominal and thoracic organs, including the liver, pancreas, stomach, spleen, and intestines, were excised to improve visualization of the aorta. The heart was held near its base to avoid compressing the ventricular chambers, and the aorta was dissected using fine microscissors and micro-forceps by first detaching it cranially at the carotid and brachiocephalic arteries, then separating it from the spine dorsally and esophagus ventrally and tracing it caudally to the iliac bifurcation. Perivascular adipose tissue was removed with fine micro-scissors to obtain a clean, intact aorta for downstream analyses.

#### Extraction of RNA from LIMA and Aorta

RNAs were extracted from LIMA and aorta according to manufacturer protocol (Direct-zol RNA MicroPrep w/TriREagent, cat# R2061-A, ZYMO RESEARCH). In Brief, cleaned aorta and LIMA were lysed in 800 and 400 µl of TRI Reagent®, respectively and mix thoroughly. Then RNAs were centrifuged and the supernatant was transferred into a new nuclease-free tube. The RNAs were purified at room temperature. An equal volume of absolute ethanol was added to samples lysed in TRI and mixed thoroughly. The mixtures were transferred into a Zymo-Spin™ IC Column2 in collection tubes and centrifuged. The columns were transferred into new collection tubes, and the flowthroughs were discarded. A400 µl RNA Wash Buffer was to the columns and the columns were centrifuged. A mixture of DNase I and DNA Digestion Buffer were mixed and added to the column matrix and was incubated at room temperature (20-30°C) for 15 minutes. Then a 400 µl Direct-zol™ RNA was added to the columns and centrifuged. RNA wash buffer (700 µl) was added to the columns, and the columns were centrifuged for 1 minute to ensure complete removal of the wash buffer. The columns were transferred carefully into an RNase-free tube and 7 and 10µl of DNase/RNase-Free water were added directly to the column matrix elute RNA from LIMA and aorta respectively, and then column were centrifuged.

#### Library Preparation

Poly(A)-enriched mRNA libraries were prepared from total RNA isolated from each whole tissue sample by Novogene using standard Illumina-compatible protocols. RNA quality and library integrity were assessed prior to sequencing. Libraries were prepared individually for each sample and sequenced on an Illumina high-throughput platform using a paired-end 150 bp sequencing strategy.

#### RNA-seq read alignment and quantification

Raw paired-end RNA-seq reads were processed and quantified using Salmon (v1.10.2) in quasi-mapping mode [23]. Transcript-level indexing was performed against the mouse reference transcriptome (GRCm39) using default Salmon indexing parameters. Reads were aligned and quantified against the indexed reference using Salmon’s lightweight alignment strategy, which accounts for sequence-specific and GC-content biases. Transcript-level abundance estimates were summarized to gene-level counts for downstream analyses using standard annotation-based aggregation.

#### Unsupervised analyses and sample-level comparisons

Unsupervised hierarchical clustering was performed using the most variable genes across samples, with distances computed using Pearson correlation and clustering performed using complete linkage. Pairwise Pearson correlation analysis was used to assess concordance among biological replicates within each vessel type. Principal component analysis (PCA) was conducted on normalized gene expression data to visualize global transcriptional differences between LIMA and aorta samples.

#### Differential expression and pathway-level transcriptomic analysis

Gene-level expression data were imported into R and analysed using the limma [24] framework. Expression values were normalized and transformed using the limma-voom method [25] to model the mean–variance relationship inherent to RNA-seq count data. Differential gene expression analysis was performed by fitting linear models comparing LIMA and aorta samples from Apoe / mice. Pathway-level analyses were performed using Gene Set Enrichment Analysis (GSEA) on ranked gene lists derived from limma differential expression statistics, using MSigDB’s curated C2 canonical pathway gene sets [26–28]. Statistical significance for differential gene expression and pathway-level analyses was assessed using empirical Bayes moderation and permutation-based testing, respectively, with multiple-testing correction performed using the Benjamini–Hochberg method. Genes with log fold change ≥ 0 and adjusted p ≤ 0.05, and pathways with FDR-adjusted q values ≤ 0.05, were considered significant.

To assess relative pathway activity at the sample level, single-sample Gene Set Enrichment Analysis (ssGSEA) was performed using predefined gene sets associated with arterial identity, oxidative stress, macrophage recruitment, inflammation, and plaque formation [27, 29, 30]. For cross-species analyses, mouse and human datasets were compared at the pathway level using ortholog-mapped gene sets. Pathway enrichment scores derived from mouse and human datasets were correlated using Spearman rank correlation to assess concordance in pathway regulation between species.

## RESULTS

### Anatomical validation of mouse internal mammary arteries

Because LIMA has never been isolated or anatomically validated in mice, our first objective was to definitively locate and characterize the murine internal mammary arteries. We selected the Apoe / model for these studies because our overarching goal was to compare the molecular signatures of the aorta and LIMA in a physiologically relevant model of atherosclerosis. Prior micro–computed tomography studies by Casteleyn et al. demonstrated that the vascular branching pattern of the aortic arch is highly conserved between mice and humans, with the brachiocephalic trunk, left common carotid artery, and left subclavian artery arising sequentially from the arch in both species [22]. Their 3D reconstructions also revealed that, in both mice and humans, the initial segment of the aorta, including the ascending aorta, aortic arch, and proximal descending aorta, forms a similar sigmoidal curvature [22].

In humans, angiographic studies show that the LIMA originates from the left subclavian artery and courses caudally along the inner thoracic wall, parallel and medial to the sternum. It lies on the posterior surface of the costal cartilages, accompanied by small veins, and gives rise to anterior intercostal branches that supply the thoracic wall. Guided by this conserved anatomy, we hypothesized that the murine LIMA follows a comparable trajectory. However, it’s extremely small caliber in mice (approximately 100–200 µm) necessitated the use of microsurgical tools and high-magnification dissection to visualize and isolate the vessel.

To locate LIMA, after skin and abdominal cavity opening, the diaphragm was exposed, and the ribcage was cut on both lateral sides **(Figure 1A).** Prior to perfusion, the thoracic wall was opened longitudinally on both sides and the ribcage was flipped towards the cranial region **(Figure 1B)**. LIMA and RIMA were located as tiny arteries 2.5 mm lateral to sternum **(Figure 1C)**. **Fig. 1D** shows LIMA pulled with a microdissection forceps. LIMA and RIMA are anatomically attached to and branch off from the inferior aspect of the left and right subclavian arteries, respectively [1, 2, 4, 5], before both descend along the anterior chest wall. As shown in **Fig. 1C**, Both LIMA and RIMA run along the inner surface of the anterior chest wall, and both run parallel to the sternum, along the posterior surface of the thoracic wall behind the sternal ends of the costal cartilages.

**Figure 1.**
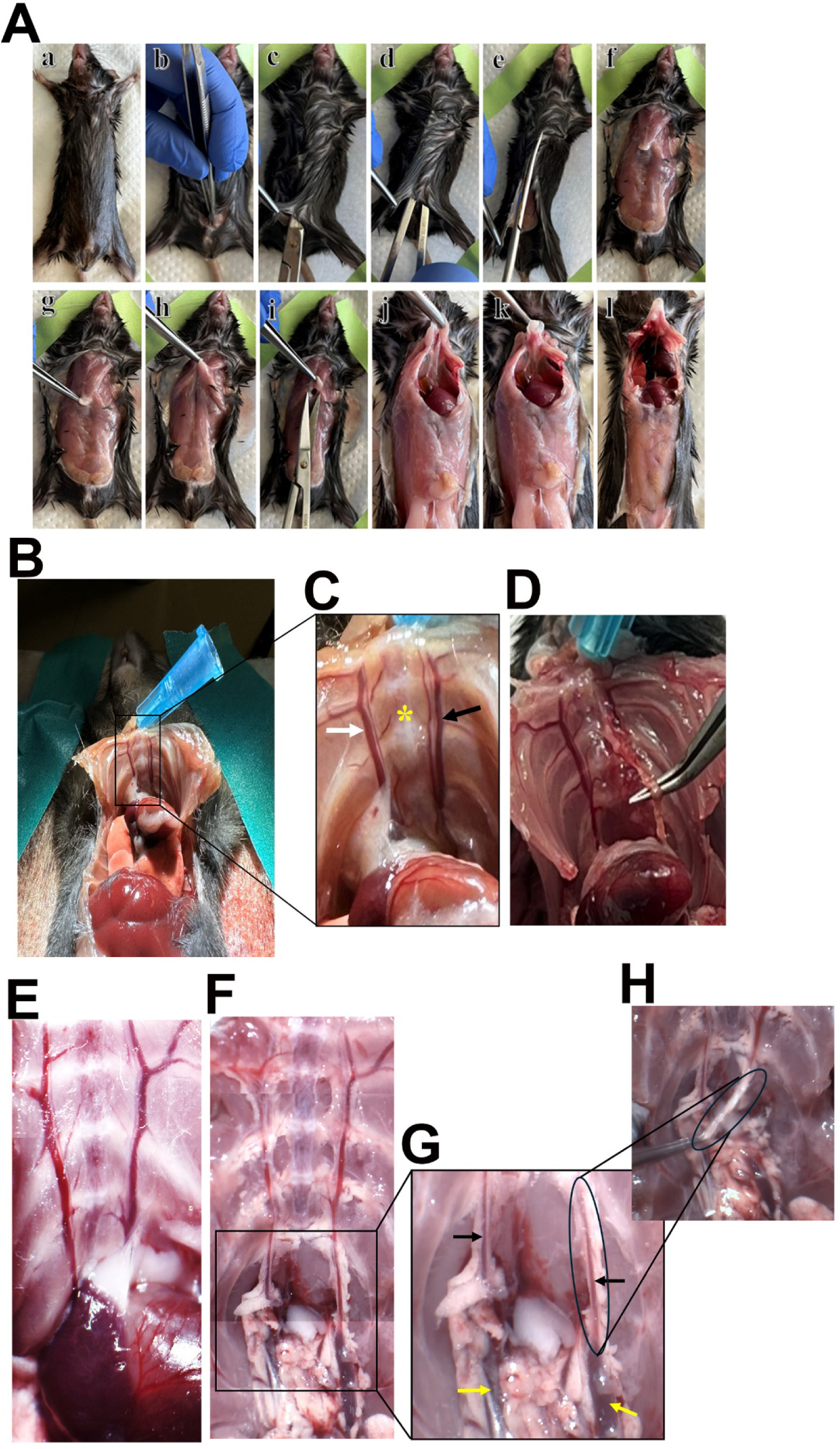
Dissection of Apoe-/- mouse to isolate IMA and Aorta. The mouse was anesthetized using CO2 in an induction chamber until the absence of pedal reflex was confirmed. Dissection was performed under aseptic conditions. Upper panel shows (A, B, C, D, E, F) (from left to right), the mouse was positioned supine on a dissection board and secured with colored tab through the limbs. An incision was made through the abdominal skin using sterile scissors, starting from the lower abdomen and extending to the thoracic cavity. The skin was gently retracted laterally to expose the abdominal wall muscles. Lower panel shows (G, H, I, J, K, L) (from left to right), an incision was made below the xiphoid process (subxiphoid incision) and then cuts through the diaphragm muscle to access the thoracic cavity of Apoe-/-. The diaphragm was carefully incised to access the thoracic cavity, revealing the organs including heart and lungs **Dissection of the internal thoracic cavity to expose LIMA.** (B) To visualize and isolate LIMA and RIMA, the inner wall of thoracic cavity was carefully dissected, and image was taken with high resolution camera to show LIMA and RIMA. (C) Both LIMA (black arrow) and RIMA (white arrow) run along the inner surface of the anterior chest wall, and both are running parallel to the sternum (yellow star), along the posterior surface of the thoracic wall behind the sternal ends of the costal cartilages. (D) Notably, LIMA is not adherent to the thoracic wall immediately upon branching from the subclavian artery, allowing for easier removal of surrounding fascia and connective tissue through this natural plane of separation. (E) The internal thoracic wall of Apoe-/- mouse post-dissection and prior to perfusion showing LIMA and RIMA originate from subclavian arteries that are branching off the aortic arch. (F) Following dissection and removal of the heart to expose the aortic arch and its three major arteries coming form the aorta shows that LIMA and RIMA originate form left subclavian artery and right subclavian artery, respectively. (G) Closer examination reveals that LIMA (right black arrow) and RIMA (left black arrow) branches off from the inferior aspect of the left subclavian artery (right yellow arrow) and the right subclavian artery (left yellow arrow) and then both are descending along the anterior chest wall. (H) Pulling out LIMA confirms anatomical attachment to subclavian artery.

Under the dissection microscope, the heart was localized adjacent to the thoracic wall with embedded LIMA and right internal mammary artery (RIMA) **(Figure 1E)**. It was gently retracted downward to expose the aortic arch along with its main branched arteries; brachiocephalic, carotid and subclavian artery, and all the surrounding adipose tissue was removed until the subclavian arteries are clearly visible **(Figure 1F)**. These arteries are the final major branches directly originating from the aortic arch. Closer examination revealed that LIMA and RIMA branch off from the inferior aspect of the left subclavian artery and the right subclavian artery and then both are descending along the anterior chest wall; pulling out LIMA confirms anatomical attachment to subclavian artery **(Figure 1G and H)**.

### Microsurgical isolation of mouse LIMA

The ribcage was fully separated off the mice and placed on a flat surface under the dissection microscope. It helped not only manage the isolation of LIMA but also to clean and remove the adipose and connective tissues using a spring scissor. LIMA was now not adherent to the thoracic wall. Isolated Pre-cleaned RIMA **(left side, Figure 2A)** and LIMA **(Right side; Figure 2A)**, surrounded by perivascular fat (PVF), were isolated and completely separated from apoe^-/-^chest wall. Isolated cleaned RIMA **(left side, Figure 2A)** and LIMA **(right side, Figure 2A)** after removal of fat and perivascular adipose tissue. **Fig. 2** shows plaque filled aorta from apoe^-/-^mouse with PVAT removed. **Supplementary Fig. 1** provides step-by-step methods of dissection and isolation of LIMA from mouse.

**Figure 2.**
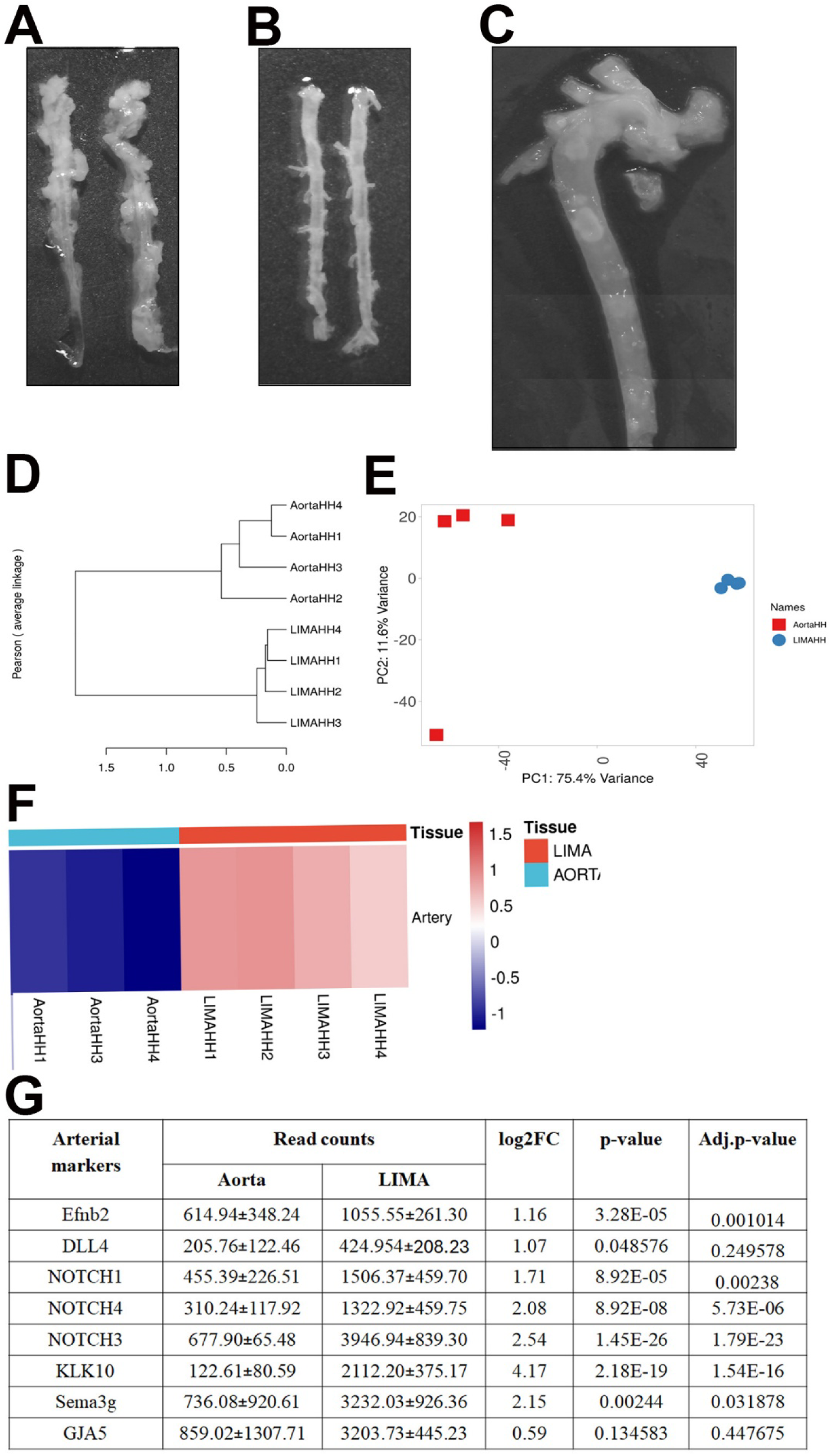
Isolating LIMA and RIMA off Apoe^-/-^ thoracic wall. (A) Isolated RIMA and LIMA before fat removal. (B) LIMA and RIMA after the removal of perivascular adipose tissue. (C) Aorta from apoe-/- mouse showing plaque build ups in the arch and branched arteries **Characterization of LIMA.** (D) Analysis of 4 LIMA and 4 aorta samples based on normalized expression values of 1541 genes. Hierarchical clustering shows the Pearson rank correlation between all samples. (E) Principal component analysis (PCA) plot showing the distribution of LIMA and aorta sample groups. (F) Heatmap showing z-scores for arterial gene modules across aorta (blue) and LIMA (red) samples. (G) Comparison of the read counts of arterial genes in aorta and LIMA of mice.

### Characterization of LIMA

We next performed transcriptomic characterization of the isolated LIMA and aorta to study the LIMA in mice. Unsupervised hierarchical clustering of the most variable genes (n = 1,247) revealed a distinct grouping of LIMA and aorta samples, indicating distinct transcriptional profiles between the two arterial tissues (Figure 2D). Pairwise Pearson correlation analysis further demonstrated high concordance among biological replicates within each vessel type. Principal component analysis (PCA) showed robust separation of LIMA and aorta along the first principal component (PC1, 75.4% variance explained), while PC2 (11.6% variance explained) captured variability among biological replicates (Figure 2E).

To assess arterial identity, we performed gene-set (module) scoring using a curated panel of canonical arterial markers. This gene set included EFNB2 (Ephrin-B2), a well-established arterial endothelial marker [31]; DLL4, a Notch ligand selectively expressed in arterial endothelial cells [32]; and the arterial regulators *NOTCH1*and *NOTCH2*, which are essential for arterial development and maintenance [33, 34]. We also incorporated *KLK10, NOTCH3, SEMA3G*, and *GJA5*, all of which are preferentially enriched in arterial cell populations [35–38]. Arterial marker expression was robust in both aorta and LIMA samples (Figure 2F–G), consistent with arterial identity. Notably, several arterial markers exhibited higher expression in LIMA compared with the aorta. In contrast, canonical venous (*NR2F2*) and lymphatic (*PROX1*, *LYVE1*) markers were expressed at substantially lower levels relative to arterial markers, consistent with an arterial transcriptional signature in both tissues.

### Transcriptomic differences in LIMA and aorta of apoe^-/-^ mice

To characterize transcriptional differences between the two arteries, we performed differential gene expression analysis comparing LIMA and aorta from Apoe / mice, revealing distinct vessel-specific transcriptional programs **(Figure 3A)**. The aorta exhibited significantly higher expression of genes associated with extracellular matrix remodelling, inflammation, and vascular pathology. These include *SFRP4*, a modulator of Wnt signaling linked to metabolic inflammation and vascular calcification. Similarly, multiple extracellular matrix–related genes, including *FMOD*, *COL5A1*, and *COL4A1*, were differentially expressed and are involved in collagen organization, fibrous cap structure, and basement membrane integrity. Inflammatory mediators such as *CCL8* is a chemokine involved in monocyte and macrophage recruitment, and *THY1* participates in leukocyte-endothelial interactions and inflammatory activation. Additionally, *ITIH4*, an acute-phase protein associated with systemic inflammation and cardiovascular risk, was differentially expressed in aorta with high significance. Together, these genes reflect coordinated differences in inflammatory, immune, and extracellular matrix pathways between the aorta and LIMA.

**Figure 3.**
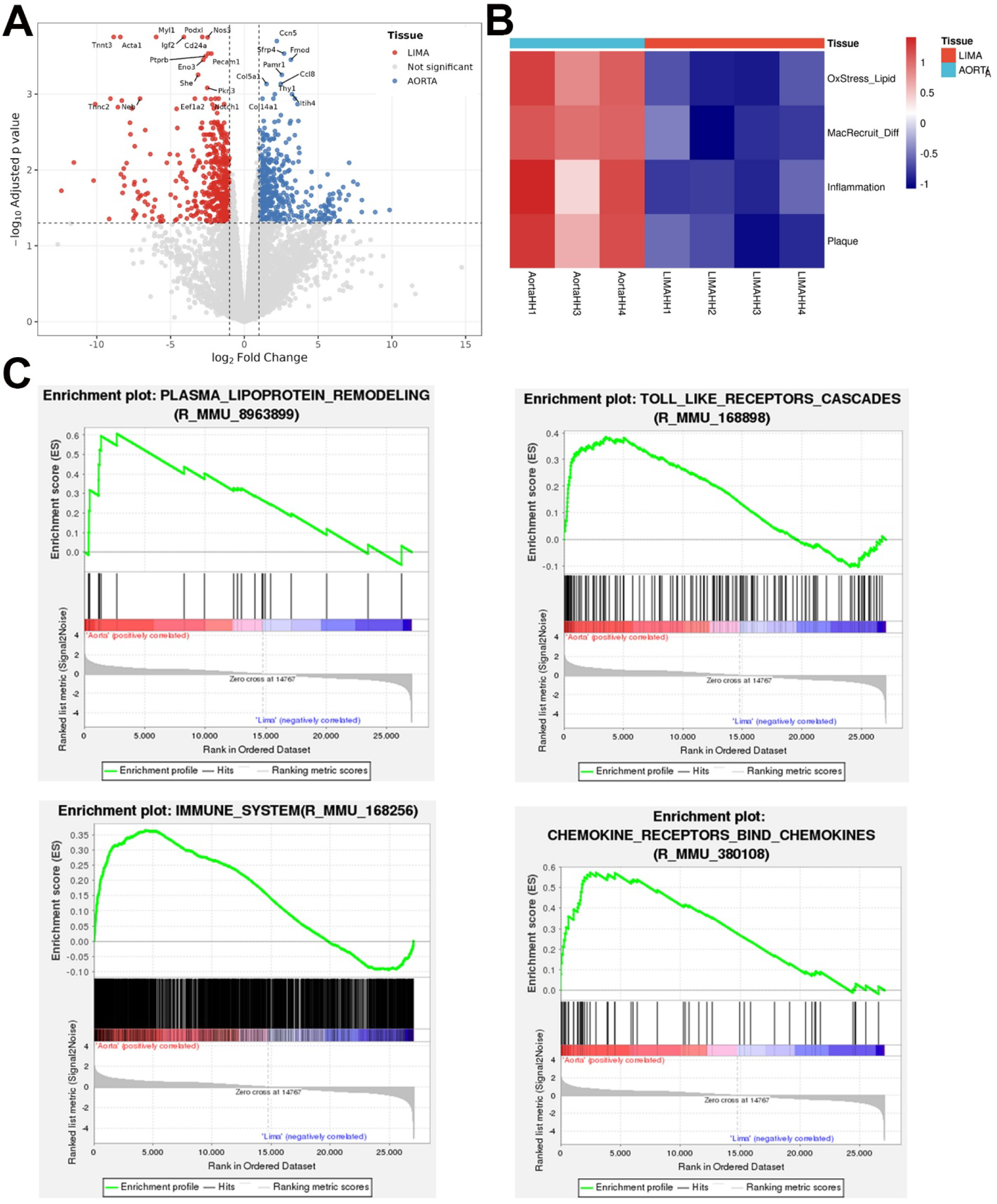
Comparison of LIMA and aorta. (A) Volcano plot shows differential gene expression between LIMA and aorta. Labeled genes denote the top significantly differentially expressed transcripts by adjusted p-value and log fold change. Red and blue dots represent genes significantly upregulated in LIMA and aorta, respectively, and grey dots denote genes with no significant change. (B) Heatmap of z-scores for atherosclerosis-related gene modules across AORTA (blue) and LIMA (red) samples. (C) GSEA pathway enrichment plots for aorta samples showing enrichment of pathways including Toll-like receptor signaling, plasma lipoprotein remodeling, immune-related pathways, and chemokine receptor signaling. The color scale indicates positively correlated genes (red) and negatively correlated genes (blue), with normalized enrichment scores (NES) and false discovery rate q values (FDR q) shown.

To further evaluate pathways relevant to plaque development, we generated heatmaps using genes associated with oxidative stress, macrophage recruitment, inflammation, and plaque formation. All genes within these atherogenic pathways exhibited markedly lower expression in LIMA compared with the aorta **(Figure 3B)**.

To further investigate biological pathways relevant to atherogenesis, we assessed the relative enrichment of pathways associated with oxidative stress, macrophage recruitment, inflammation, and plaque formation. Compared with the aorta, LIMA exhibited reduced activation of these atherogenic pathways, reflecting a markedly attenuated pro-atherogenic transcriptional profile **(Figure 3B).**

Gene Set Enrichment Analysis (GSEA) corroborated these findings, demonstrating significant enrichment of key pathological pathways of atherosclerosis in the aorta relative to LIMA **(Figure 3C)**. GSEA shows the activation of lipoprotein remodeling, Toll-like receptor signaling, and immune-system activation, and chemokine-receptor pathways, which are foundational to plaque initiation, progression and rupture **(Figure 3C)**. Also, multiple biological processes that shape plaque composition, stability, and thrombosis risk and amplify inflammation, including neutrophil degranulation, platelet activation, matrix metalloproteinase signaling, and fibrin clot formation, were activated in the aorta relative to LIMA (**Supplementary Figure 2**). Together, these data indicate that the aorta displays strong transcriptional activation of inflammatory, immune, and thrombogenic pathways, whereas the LIMA exhibits a substantially muted atherogenic signature.

### Comparison of human and mouse LIMA

To gather the clinical evidence of genes expressed in LIMA and atherosclerotic arteries in humans, we utilized data collected by Sulkava et al [39], where they performed transcriptomic analyses on human artery samples collected in the Tampere Vascular Study. The analysis was performed on atherosclerotic human artery samples from the aorta, carotid, and femoral vessels, and LIMA. To determine whether the transcriptional differences observed between mouse LIMA and aorta are conserved in humans, we performed cross-species pathway enrichment analysis using matched datasets from human LIMA and atherosclerotic aorta. Pathway-level comparison revealed a strong positive correlation in enrichment patterns between human and mouse vessels. Specifically, a Spearman correlation analysis demonstrated a significant concordance in the direction and magnitude of pathway regulation across species **(Figure 4)**. This cross-species alignment indicates that key biological programs distinguishing LIMA from the aorta, particularly those related to inflammation, immune activation, and extracellular matrix remodeling, are preserved in both humans and mice, supporting the translational relevance of the mouse LIMA model.

**Figure 4.**
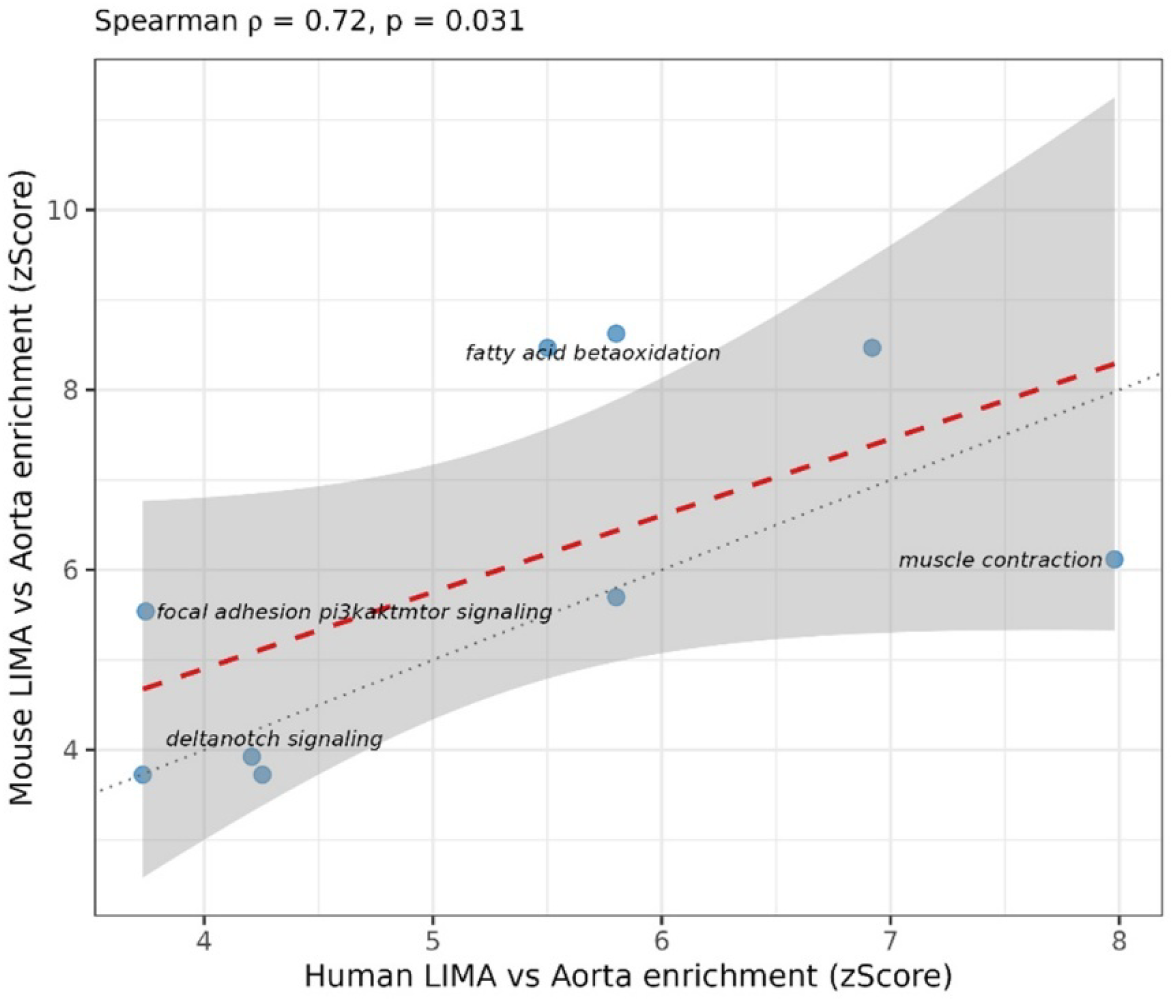
Comparison of human and mouse LIMA. Cross-species Spearman correlation (ρ = 0.72, p = 0.031) of pathway enrichment between human and mouse LIMA versus AORTA.

To further confirm, we extrapolated the data from Sulkava et al. study for the genes associated with the pathogenesis of atherosclerosis in the aorta vs LIMA and compared then to that in the apoe^-/-^ mice. Excitingly, all these genes associated with atherosclerosis showed similar trend in both human and mouse samples, i.e. higher in aorta compared to LIMA **(Supplementary Figure 3)**. The most biologically relevant atherosclerosis-related genes across the datasets included classical regulators of lipid handling (ApoE, ABCA1, CD36) [40–42], oxidative stress mediators (CYBB, HMOX1, GPX1) [43–45], key endothelial adhesion and leukocyte recruitment molecules (VCAM1, SELE, PECAM1, CXCR4, ITGB2/ITGAM/ITGAL, CD44) [46–48], major inflammatory cytokines and receptors (IL6, IL1B, CCL2, CXCL10, TLR4, NLRP3, PLA2G7) [49–53], macrophage markers (CD68) [54], and central drivers of plaque remodeling and calcification (SPP1, TIMP1, FN1, RUNX2, KLF4) [55–59]. The concordance of these gene expression patterns across species indicates that the molecular distinction between the aorta and LIMA is conserved in humans and mice, with the aorta uniformly exhibiting a transcriptional profile enriched for genes linked to atherogenesis.

## DISCUSSION

In this study, we uncover a deeply conserved molecular architecture that distinguishes atheroprone from atheroresistant vascular beds and introduce a technically enabling approach for isolating the mouse LIMA, an arterial segment previously inaccessible for mechanistic investigation. By integrating transcriptomic datasets from human arteries with newly generated profiles from Apoe / mouse aorta and LIMA, we demonstrate that regional vascular susceptibility to atherosclerosis is governed by transcriptional programs that are remarkably preserved across species. Interestingly, the conserved suppression of inflammatory and immune pathways in the mouse and human LIMA strongly supports the biological relevance of our mouse LIMA isolation method and validates its utility as a translational model for investigating molecular determinants of vascular protection.

We confirmed that the murine LIMA closely mirrors the anatomical organization of the human internal mammary artery, including its origin from the subclavian artery and its trajectory along the posterior sternum [60]. The ability to reproducibly expose, validate, and isolate this small-caliber artery is a strength of the present study, and it allowed us to generate the first transcriptomic atlas of the mouse LIMA. A central finding of our work is that the aorta, irrespective of species, exhibits a consistent upregulation of canonical atherogenic pathways, including lipid handling and retention, endothelial activation, leukocyte recruitment, and inflammatory signaling. In contrast, the LIMA retains a stable, quiescent phenotype enriched for pathways that support extracellular matrix remodeling, endothelial integrity, and suppression of inflammatory activation. This dichotomy mirrors clinical observations in which the LIMA remains protected even in the context of systemic cardiovascular risk factors and advanced aortic disease. The strong cross-species correlation in pathway enrichment provides compelling evidence that these protective and pathogenic signatures are evolutionarily encoded rather than species-specific phenomena.

Importantly, our ability to isolate and molecularly characterize mouse LIMA represents a major methodological advance. The LIMA has long been recognized in humans as a uniquely athero-resistant artery, yet its mechanistic protective features have remained largely unexplored because of the absence of an analogous small-animal model. Our new isolation technique closes this gap, enabling controlled experimental manipulation of an atheroresistant vascular bed in a genetically tractable system. This innovation allows for direct hypothesis testing, such as perturbation of endothelial signaling, extracellular matrix components, or anti-inflammatory regulators, that were previously not possible.

The translational implications of these findings are substantial. First, the conserved transcriptional differences uncovered here provide a molecular blueprint for understanding why certain arteries resist plaque formation despite systemic atherogenic stress. Second, our approach establishes the mouse LIMA as a new experimental platform for discovering protective genes, pathways, and therapeutic targets that may be harnessed to “LIMA-ize” atheroprone arteries. Finally, the concordance between human and mouse arterial biology validates the use of this model for probing regional vascular susceptibility and for testing interventions aimed at enhancing arterial resilience.

Collectively, our study provides foundational insight into the conserved biology of atherosclerosis susceptibility and introduces a transformative methodological resource for the field. By pairing mechanistic discovery with a newly accessible murine vascular bed, this work opens the door to targeted strategies that leverage nature’s own blueprint for atheroprotection.

## ACKNOWLEDGEMENTS

This work was supported by NIH/NIDDK grant R01DK101711 (VP), R01HL140836 (VP), funds from Osteopathic Heritage Foundation’s Vision 2020 to Heritage College of Osteopathic Medicine at Ohio University (VP).

## AUTHOR CONTRIBUTIONS

H.H., P.S., K.W.J., K.P., L.Z. contributed to study design, performing experiments, analysis of results, and manuscript preparation. B.B. contributed to study design, performing experiments, and data analysis. M.B. and S.K.M. contributed to data analysis. V.P. contributed to study design and oversight, performing experiments, analysis of results, and manuscript preparation.

## FIGURE LEGENDS

**Supplementary Figure 1:**
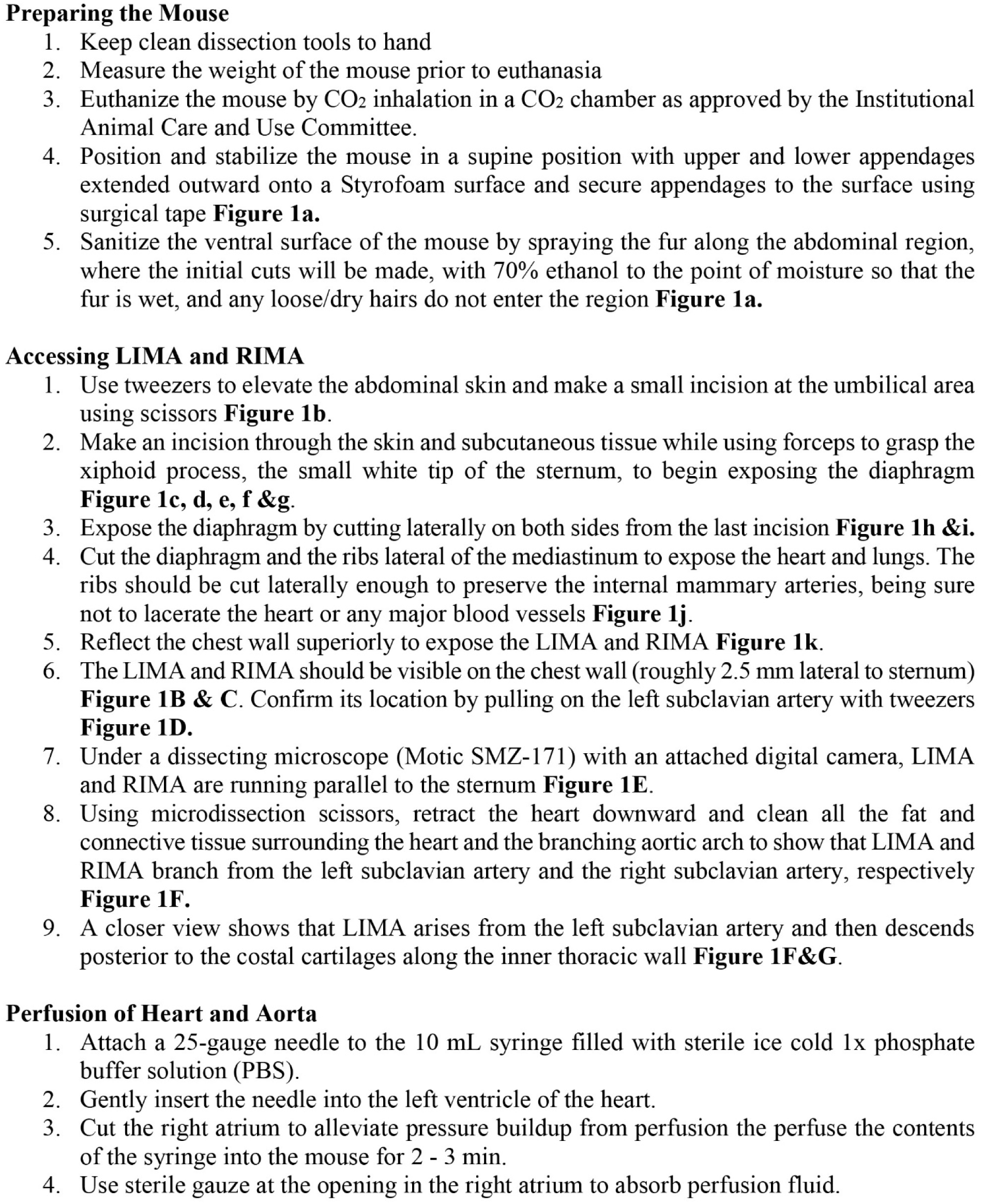
Step-by-step method of dissection and isolation of LIMA from mouse.

**Supplementary Figure 2:**
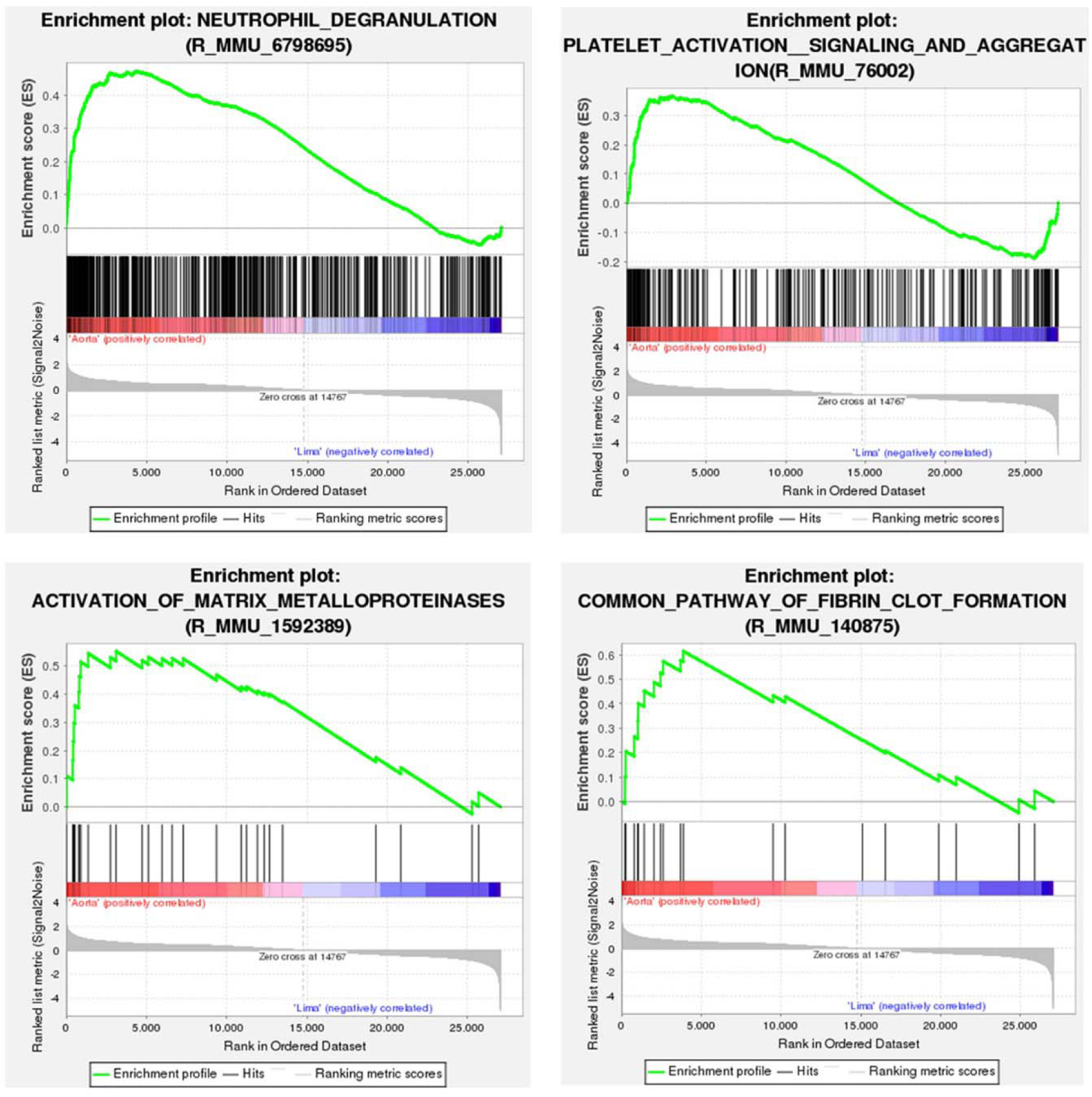
KEGG GSEA pathway-enrichment plots showing upregulated pathways include neutrophil degranulation, platelet activation, activation of matrix metalloproteinases, and fibrin clot formation. The colored horizontal bar indicates a shift from positively correlated genes (red) to negatively correlated genes (blue). The normalized enrichment scores (NES) and false discovery rate q value (FDR q) are indicated.

**Supplementary Figure 3:**
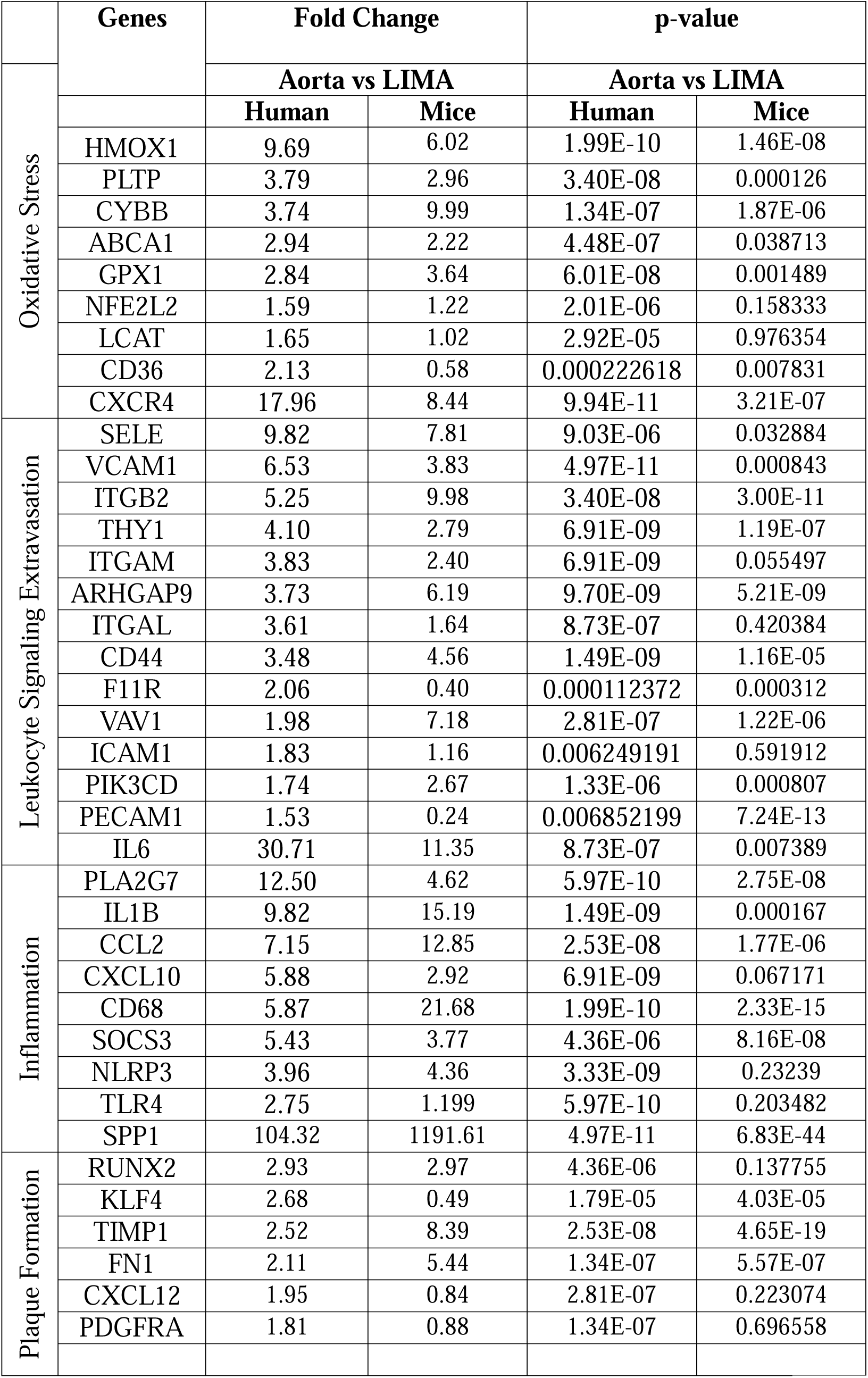
Genes associated with atherosclerosis in aorta vs LIMA of apoe^-/-^mice. The most biologically relevant atherosclerosis-related genes across pathways including oxidative stress, endothelial adhesion and leukocyte recruitment, inflammatory cytokines and receptors, and plaque formation. The data shows fold change of expression of genes in aorta vs LIMA.

